# The Whsc2/NelfA -dependent transcription complex is required for postnatal cardiac development and heart function

**DOI:** 10.1101/2025.10.24.684384

**Authors:** Saleena Alikunju, Chandni Thakkar, Aishwarya Venkatasubramanian, Zhi Yang, Andreas Ivessa, Danish Sayed

## Abstract

Cardiac malformations and ventricular remodeling due to heart diseases result in compromised cardiac function, eventually leading to heart failure. In this study, we examine the role of cardiac Wolf-Hirschhorn Syndrome candidate 2 (Whsc2), *also known as* Negative elongation factor A (NELFA), one of the genes encoded in the WHS critical region. The Wolf-Hirschhorn Syndrome is a contiguous genetic disorder due to microdeletions in the critical region, with clinical manifestations of neurological defects frequently associated with congenital malformations, including cardiac defects. NelfA has been implicated in RNA polymerase II (pol II) pausing, suggesting a role in pol II-dependent gene transcription. We previously reported an early onset of heart failure with the acute knockdown of NelfA in hearts undergoing pressure overload-induced cardiac hypertrophy. Here, we characterize a mouse model with cardiomyocyte-specific loss of NelfA function, in which these mice develop spontaneous cardiomyopathy at 2 months of age and exhibit early mortality by 3 months, suggesting a critical role for postnatal NelfA in the heart. Interactome data show that chromatin-bound NelfA interacts with proteins involved in chromatin remodeling (Trim28) and pre-mRNA processing (Adrph1l), along with expected binding partners like RNA pol II, Supt5, and other Nelf proteins. Examination of genomic occupancy of these NelfA-associated proteins in the NelfA knockout (KO) hearts reveals a disassembly of the NelfA nucleated complex at promoters of cardiac-enriched genes, including cytoskeletal and metabolic genes. This deconstruction of the NelfA-dependent complex results in the inhibited expression of these essential genes during postnatal cardiac development, leading to a cardiac contractile and metabolic crisis that precipitates dilated cardiomyopathy.

## Introduction

Wolf-Hirschhorn Syndrome candidate 2 (Whsc2), *also known as* Negative elongation factor A (NELFA), is one of the genes encoded in the WHS critical region. The Wolf-Hirschhorn Syndrome (WHS), with an incidence of 1 in 20-50,000 live births, has clinical manifestations of severe neurological defects along with congenital malformations, including cardiac defects. The microdeletions and translocations in the WHS critical region (WHSCR) on chromosome 4 result in haploinsufficiency of genes, which have been linked to variable degrees of clinical manifestations [1, 2].

Whsc2 or NelfA is a core component of the negative elongation factor complex, which has been shown to play a role in regulating RNA polymerase II (pol II) pausing, thereby regulating pol II-dependent gene transcription [3, 4]. RNA pol II pausing has been identified as the rate-limiting step in eukaryotic transcription and as a widespread mode of gene regulation, as opposed to initial reports in Drosophila, where it was thought to be restricted to heat shock proteins or genes that require rapid transcription [5-8]. Negative elongation factor (Nelf) complex, comprising four or five subunits (A to E) in coordination with DRB sensitivity-inducing factor (DSIF), has been implicated in mediating pol II pausing, which is released by P-TEFb (Cyclin T/Cdk9) [3, 4]. In vitro studies and structural examination of the paused complex established the orientation of the Nelf proteins, Supt5, and pol II within the complex, and predicted that NelfA is indispensable for assembly of the paused complex [9, 10]. While investigating active transcription in mouse hearts undergoing cardiac hypertrophy, we reported widespread Pol II pausing at gene promoters of mostly constitutively expressed essential genes. Interestingly, we observed a synchronous, incremental release of paused Pol II into productive elongation, which is commensurate with the increase in cardiac mass that occurs when mice are subjected to pressure overload [11, 12]. These results suggested that the promoter clearance of paused Pol II contributed to accommodating the increase in basic cellular functioning. Furthermore, we demonstrated that an increase in NelfA expression and promoter occupancy is necessary for the active transcription of both essential and inducible genes during cardiac hypertrophy. Acute knockdown of cardiac NelfA precipitated the early onset of cardiac failure in mice subjected to pressure overload [13].

In this study, we characterize the NelfA knockout (NelfA-KO) mouse model, which exhibits loss of postnatal NelfA function and develops spontaneous cardiomyopathy due to the disassembly of the NelfA-dependent complex at cardiac-enriched gene promoters. NelfA-KO hearts present with contractile and metabolic dysfunction, which precedes cardiac failure, the development of cardiomyopathy, and death.

## Materials and methods

### Animals

All animal procedures in this study were performed per the NIH Guidelines for the Care and Use of Laboratory Animals (National Academies Press, No. 85-23, 2011). The animals were housed in the Animal Facility at Rutgers, The State University of New Jersey, Newark, NJ, per standard procedures/protocols. The IACUC of the Rutgers-New Jersey Medical School approved all protocols. C57BL/6 mice for Sham and transverse aortic constriction (TAC) surgeries were purchased from The Jackson Laboratory. Cardiac-specific NelfA-KO mice were generated by crossing NelfA(flox/flox) mice (generated at Cyagen Biosciences) with αMHC-Cre mice. Both male and female mice were used and compared to their respective controls concerning sex and age. Anesthesia: - Mice were injected with Ketamine/Xylazine/Acepromazine (80-100mg/Kg / 5-12mg/Kg / 1-2mg/Kg) intraperitoneal. Analgesia: - Buprenorphine Ethiqa XR (3.25 mg/Kg) subcutaneously every 72 hrs post-surgery. Euthanasia: - All experimental animals (except the cohort for the survival study) undergo terminal echocardiography under anesthesia, followed by tissue extraction for biochemical and molecular analysis.

### Transverse Aortic Constriction

TAC was performed as described previously [12, 13]. A 7-0 braided polyester suture was tied around the transverse thoracic aorta against a 27-gauge needle between the innominate artery and the left common carotid artery. Control mice were subjected to a sham operation involving the same procedure minus the aortic constriction.

### Survival analysis

Cohorts of 38 NelfAKO mice (14 Male and 24 Female) and age and sex-matched Wt Cre were recruited and observed daily to record their survival. The event was entered as “1” when mice were found dead, while a “0” was entered when the mice were found healthy at the endpoint of observation. Survival analyses were performed using the Kaplan-Meier survival analysis plot in GraphPad Prism 10.0.2.

### Echocardiography

Echocardiographic measurements were performed as described previously [12, 13]. Briefly, mice were anesthetized with Ketamine (80-100 mg/kg)/Xylazine (10mg/kg) administered intraperitoneally. The tail pinch confirmed the adequacy of the anesthetic. Transthoracic echocardiography was performed using the Vevo 3100 imaging system (Visual Sonics) with an MX400 30 MHz (mouse, cardiology) scan head encapsulated transducer. Electrocardiographic electrodes were taped to the four paws, and then 1D M-mode and 2D B-mode tracings were recorded from the parasternal short-axis view at the mid-papillary muscle level. Vevo 3100 Software (Vevo Lab, version 3.2.6) was used for image capture and analysis.

### ChIP-Seq

Chromatin immunoprecipitation sequencing (ChIP-Seq) was performed, as reported earlier [14]. Hearts from 55-day-old male mice were harvested, snap-frozen in liquid nitrogen, and sent to Active Motif (Carlsbad, CA) for Supt5-, NelfE-, Trim28-, and RNA pol II-ChIP-Seq. Immunoprecipitation was performed using antibodies against the listed proteins, followed by high-throughput Illumina NextSeq 500 sequencing. Input DNA was taken before immunoprecipitation and served as a control for normalization and eliminating background.

### ChIP-Seq Data Analysis

Bioinformatics on the sequencing data was performed by Active Motif, Inc., and done as described previously [14]. The Data explanation from Active Motif includes the following: *Sequence analysis* – 75-nt sequence generated by sequencing reads was mapped to the genome using the BWA algorithm, and the information was stored in BAM format. Only reads that align with no more than two mismatches and map uniquely are used in the analysis. *Determination of Fragment Density* – Aligned reads (tags) are extended in silico at their 3′ ends to a length of 200 bp using Active Motif software, corresponding to the average fragment length in the size-selected library. To identify the density of fragments along the genome, the genome is divided into 32-nt bins, and the number of fragments in each bin is determined and stored in a BigWig file and can be visualized on genome browsers. *Peak Finding* – Genomic regions with enrichments in tag numbers are termed ‘intervals’, and defined by the chromosome number, start and end coordinate. Normalized tags were used for peak calling using the MACS 2.1.0 algorithm with a default cutoff p-value of 1e-7. Peak filtering was performed by removing false ChIP-Seq peaks as defined within the ENCODE blacklist. *Merged region analysis* – To compare peak metrics between two samples, overlapping Intervals are grouped into ‘Merged Regions’, which are defined by start coordinate of the most upstream interval and end coordinate of the most downstream interval (=union of overlapping intervals; “merged peaks”). In locations where only one sample has an interval, this interval defines Merged Region. The use of Merged Regions is necessary because the locations and lengths of intervals are rarely exactly same when comparing different samples. *Annotations* – Intervals, Merged Regions, their genomic locations along with proximities to gene annotations and other genomic features are determined and presented in Excel spreadsheets. Average and peak fragment densities within Intervals and Active Regions are compiled.

The Merged Regions provide the peak metrics for samples in all peak regions and are used to analyze ChIP enrichments in the samples. Pairwise comparisons were performed using MaxTags, followed by Log2 ratio. To calculate the maximum number of tags (reads), multiply the average value of the tag per sample by the length of the regions and divide by a constant of 224. The 224 value represents the average length of the sequenced fragment in base pairs. As the signal is normalized in 32 bp bins along the genome, 7 bins of 32 bp length (224) are equivalent to the fragment length of 200 bp. From the sorted data, subsets were created for manuscript preparation, ensuring proper data representation and presentation.

### RNA Sequencing

Hearts from 55-day-old NelfA-KO male mice and their corresponding Wt-Cre controls were collected and sent to Novogene Corporation Inc. for RNAseq. Briefly, mRNA was purified from total RNA using poly-T oligo-attached magnetic beads. After fragmentation, the first strand cDNA was synthesized using random hexamer primers, followed by the second strand cDNA synthesis using either dTTP for a non-strand-specific library or dUTP for the strand-specific library. These libraries were ready after end repair, A-tailing, adapter ligation, size selection, amplification, and purification. The libraries were checked with Qubit and real-time PCR for quantification and bioanalyzer for size distribution detection. After library quality control, different libraries were pooled based on the effective concentration and targeted data amount and then subjected to Illumina sequencing. Sequenced reads were processed through fastp software to remove possible adapter sequences and nucleotides with poor quality to obtain clean reads. Paired-end clean reads were aligned to the reference genome using Hisat2 v2.0.5. Alignment information for each read was stored in the BAM format. The mapped reads of each sample were assembled by StringTie (v1.3.3b) in a reference-based approach. Feature Counts v1.5.0-p3 was used to count the reads mapped to each gene. The distribution of read counts in libraries was examined before and after normalization. The original read counts were normalized to account for various factors, including variations in sequencing yield between samples. These normalized read counts were used to accurately determine differentially expressed genes using the DESeq2 R package (1.20.0). Log2 fold change between the groups was calculated, and the Wald test was used to determine statistical significance (p-value). Adjusted p-values were calculated using the Benjamini-Hochberg procedure. Genes with an adjusted p-value <0.05 and log2FC of at least one are called out as differentially expressed genes to compare the groups. A heatmap for the differentially regulated genes was generated using Heatmapper [15], with average linkage for clustering.

### Genome browsers

Integrated Genome Browser (IGB) [16] and Integrated Genomic Viewer (IGV) [17] were used to visualize RNAseq data, respectively.

### Functional annotation and GO terms

Functional annotation was performed using the Database for Annotation, Visualization, and Integrated Discovery (DAVID) algorithm [18].

### Western Blotting

Protein lysates were fractionated into cellular compartments as per the manufacturer’s protocol (ThermoFisher Scientific; cat # 87790). Protein was quantified by BCA protein assay kit (Thermo Scientific), separated on gradient (4%–12% gel) XT gels (Biorad), and transferred to 0.2 μm nitrocellulose membrane. Membranes were probed with primary antibodies for the indicated proteins. The Western blot signals were detected by the Odyssey imaging system (LI-COR).

### Quantitative PCR

Total RNA was isolated from cardiomyocytes using TRIzol reagent (Life Technologies) and reverse-transcribed to cDNA using a High-Capacity Reverse Transcription Kit (Thermo Fisher Scientific), per manufacturer’s protocol. The cDNA was used for qPCR with an Applied Biosystems 7500 thermocycler using Taqman gene expression assays. 18S was used as an internal control for normalization.

### Histological analysis

Hearts were fixed with 10% formalin and embedded in paraffin. Paraffin-embedded tissue slices (5 um) on slides were deparaffinized, rehydrated, and stained with Picric Acid Sirius Red (fibrosis), Wheat Germ Agglutinin (cell size), and Hematoxylin & Eosin (tissue structure and cell morphology) stains. Tissue slices were washed, dehydrated, and mounted in a permanent mounting medium.

### Statistics

ChIP-Seq statistics (provided by Active Motif), RNA-Seq statistics (provided by Novogene), and metabolomics statistics (provided by Creative Proteomics) are detailed in the respective sections. Calculation of significance between 2 groups was performed using an unpaired, 2-tailed Student’s t-test. All experiments were conducted with a minimum of n=3 and presented as average with SEM; p<0.05 was considered significant.

## Results

### NelfA is required for postnatal cardiac development and function

To determine the transcriptional role of NelfA in the heart, we generated conditional cardiac-specific NelfA knockout (NelfAKO) mice. The RNA polymerase II binding domain of NelfA, encoded by exons 4-7, was flanked by LoxP sites. Homozygous LoxP (LoxP+/+) mice were mated with αMHC-Cre mice for late fetal knockout of NelfA in the heart (**Fig. 1A** and **1B**). Interestingly, these mice begin to exhibit signs of spontaneous heart dysfunction and cardiomyopathy development ∼60 days after birth. Functional characterization of these hearts (55-60days) shows a significant increase in heart weight (**Fig. 1C**), a decrease in percent ejection fraction and fractional shortening (**Fig. 1D**), indicating cardiac dysfunction, an increase in systolic and diastolic left ventricular dimensions (**Fig. 1E**), along with an increase in left ventricular volume after systole and diastole (**Fig. 1F**), suggesting chamber dilatation and onset of dilated cardiomyopathy. These NelfAKO mice exhibit 100% mortality by the age of 110-140 days (**Fig. 1G**), characterized by dilated, thinned-walled ventricles compared to their respective wild-type (Wt-Cre) hearts (**Fig. 1G**). Morphological analysis reveals an increase in individual cardiomyocyte size (**Fig. 1I**), an increase in cells undergoing apoptosis (**Fig. 1J**), and an increase in perivascular and interstitial fibrosis (**Fig. 1K**). These results indicate that NelfA is required for postnatal cardiac development and function, and loss of NelfA function results in spontaneous cardiac dysfunction, heart failure, and death.

**Fig. 1.**
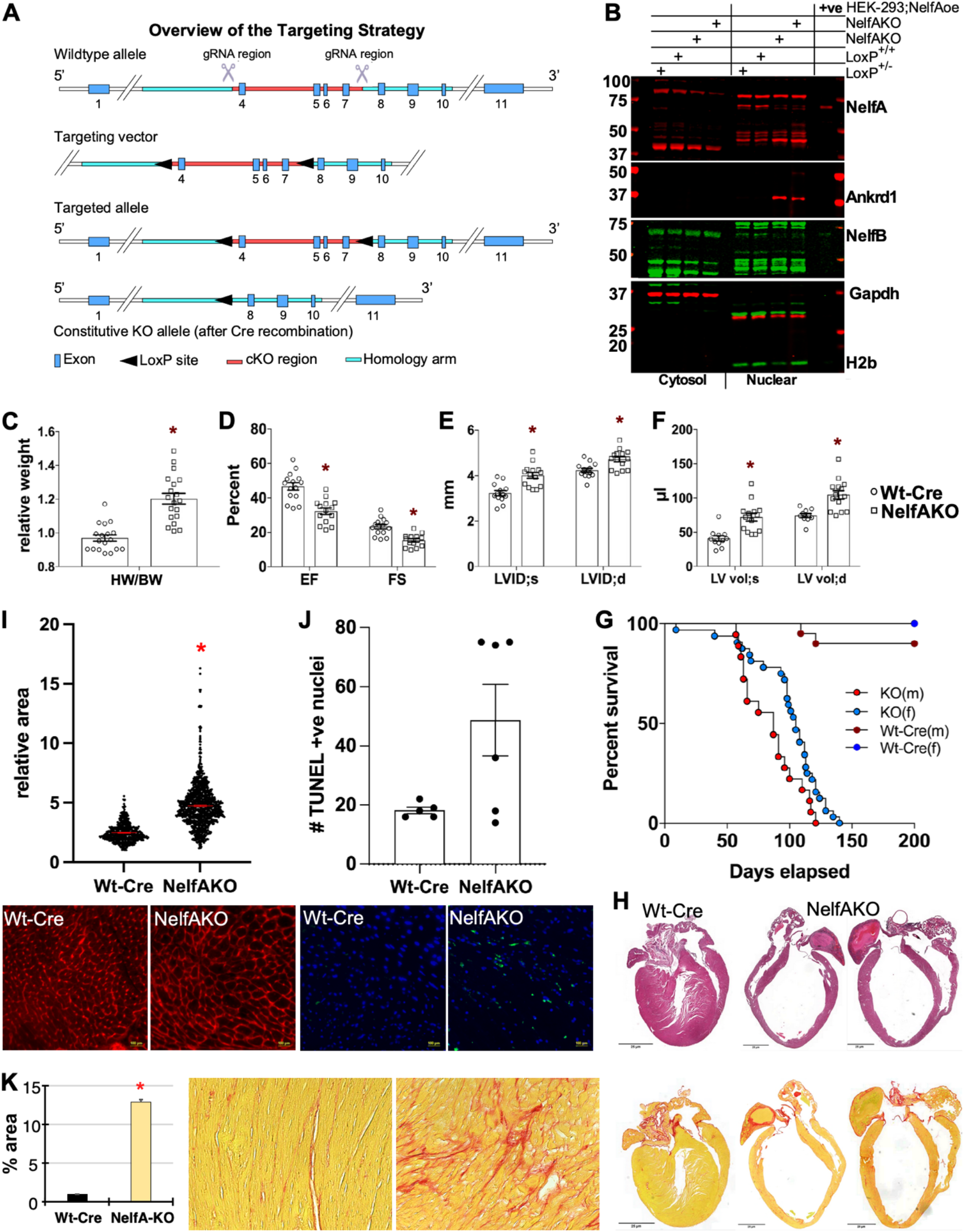
Generation and physical, morphological, and functional characterization of NelfAKO mice. **A**. Schematic shows the overview of the targeting strategy for the generation of conditional cardiac-specific NelfAKO mice. Homozygous floxed (LoxP) mice were mated with αMHC-Cre mice for late fetal knockout of NelfA in the heart. **B**. Western blot analysis shows the expression levels of the indicated proteins, along with their subcellular localization. Ankrd1 (Carp) is shown as a marker for cardiac hypertrophy, NelfB is shown for specificity, Gapdh and H2B are shown for compartment specificity and loading control. The last lane serves as a positive control, consisting of protein lysate from HEK293 cells transfected with a plasmid expressing NelfA cDNA under the CMV promoter. **C–F**. The graphs represent heart weight per body weight (HW/BW), percent ejection fraction (%EF) and percent fractional shortening (%FS), left ventricular internal dimensions during systole and diastole (LVID;s, LVID;d), and left ventricular volume with systole and diastole (LV Vol;s and LV Vol;d), respectively in Wt-Cre and NelfAKO mice, as indicated and measured by echocardiography. **G**. Survival curve for Wt-Cre and NelfAKO mice (m for males and f for females) generated on Prism 10 on cohorts (n = 24 females and n = 14 males NelfAKO mice with their respective Wt-Cre controls. **H**. H & E staining of longitudinal sections of hearts from Wt-Cre and NelfAKO mice showing chambers, wall thickness, and fibrosis. **I**. The graph represents the cross-sectional area of individual cardiomyocytes in sections from hearts of Wt-Cre and NelfAKO mice. Representative images of the WGA staining are shown. **J**. The graph represents the number of TUNEL-positive nuclei from the hearts of Wt-Cre and NelfAKO mice. Representative images of the TUNEL staining are shown. **K**. The graph describes the mean fibrotic area in sections from hearts of Wt-Cre and NelfAKO mice, as measured using Image J. Representative images of PASR staining in Wt-Cre and NelfAKO mice showing cardiac fibrosis. For all graphs, error bars represent SEM. * indicates p < 0.05 compared to the respective Wt-Cre group, n > 5 for each group, unless indicated otherwise.

### NelfA-dependent complex in the heart

To decode the chromatin-bound NelfA complex, we performed interactome profiling using Rapid Immunoprecipitation Mass Spectrometry of Endogenous proteins (RIME) in sham and TAC-induced hypertrophied hearts. Along with the Nelf subunits (NelfB, NelfC/D, NelfE), Supt5 and RNA polymerase II, other associated proteins include Trim28, Adprhl1, Numa1, and Hist1h1e (**Fig. 2**). In accordance with our previous observations that NelfA is associated with active gene promoters, these results reveal NelfA-dependent complex occupancy on cardiac gene promoters in both sham and TAC hearts, indicating that the Nelf complex does not dissociate from the chromatin during cardiac hypertrophy, which is associated with a compensatory increase in gene expression. RIME data further suggest that the NelfA-dependent complex is involved in more than just RNA pol II pause-release and might be involved in chromatin remodeling and nascent transcript processing.

**Fig. 2.**
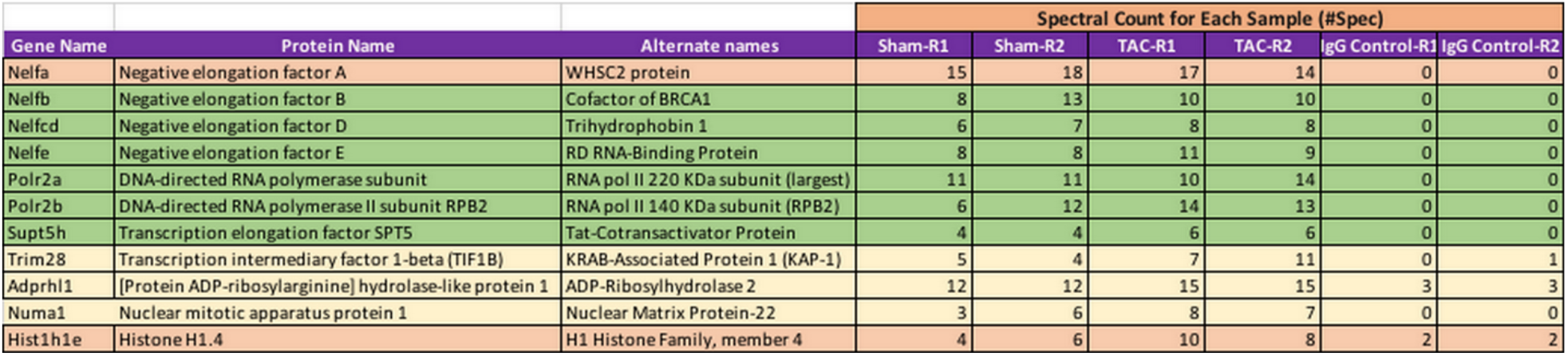
NelfA interactome in the heart. The table shows the spectral counts for proteins associated with chromatin-bound NelfA as detected by Rapid Immunoprecipitation Mass Spectrometry of Endogenous proteins (RIME). R1 and R2 represent two sets (3-4 hearts in each set) run as independent samples. IgG is the control. The cutoff for spectral counts was maintained at 5 in sham and TAC hearts.

### Loss of NelfA disrupts the chromatin-bound NelfA-dependent complex at cardiac-enriched gene promoters

To examine the status of chromatin association of Supt5, NelfE, Trim28, and RNA pol II in NelfAKO and Wt-Cre hearts, we performed ChIP-Seq in 55-day-old male mouse hearts. Fig. 3A represents the heatmaps of the merged regions of Supt5, NelfE, Trim28, and RNA pol II, showing the occupancy status on the cardiac genome and distribution across promoters and in-gene regions (**Fig. 3A**). Comparing the overall sequencing tag counts for these NelfA complex proteins in the KO and WT-Cre hearts reveals a decrease in total NelfE and Trim28 tags in KO hearts, while little or no change is observed in Supt5 and RNA pol II tag counts in the KO versus the WT-Cre hearts (**Fig. 3B**). Further, as expected, Supt5, NelfE, and Trim28 occupancy is enriched at the promoter regions, while ∼40% of RNA pol II is promoter bound in NelfAKO and Wt-Cre hearts (**Fig. 3C**). To further characterize the ChIP-seq data sets, we used the differential occupancy status of Supt5 as a measure to categorize genes. The average occupancy densities of these factors at the promoter and in-gene regions were aligned together, along with their respective inputs. The 54.22% (14498 out of 26735) of genes with average pol II density at the promoter or in-gene regions equal to or greater than 6.5 (average density of input performed with RNA pol II-ChIP-seq) were considered as expressed and included for further analysis. Furthermore, these 14498 genes were separated based on Supt5 average densities equal to or greater than 1.8 (average density of the input, as determined with Supt5-ChIP-Seq), and 49.91% (13346 out of 26735) or 92.05% (13346 out of 14498) were included in the differential analysis between NelfAKO and Wt-Cre hearts. Of the 13346 genes, 515 genes show a decrease in Supt5 promoter occupancy (2-fold difference). This set of genes also exhibits a significant decrease in overall Supt5 genomic binding, a decrease in NelfE and Trim28 genomic binding at the promoter and intragenic (in-gene) regions, a decrease in RNA polymerase II promoter occupancy, but no change in intragenic RNA polymerase II distribution and binding (**Fig. 3D** and **3I**). These results suggest that the loss of NelfA results in disrupted complex assembly at these gene promoters, likely hampering gene transcription. Conversely, only 78 genes show an increase in promoter Supt5 (Log2FC≥1), which does not correlate with similar changes in NelfE or Trim28, but an increase in promoter RNA pol II is observed (**Fig. 3E** and **3I**).

**Fig. 3.**
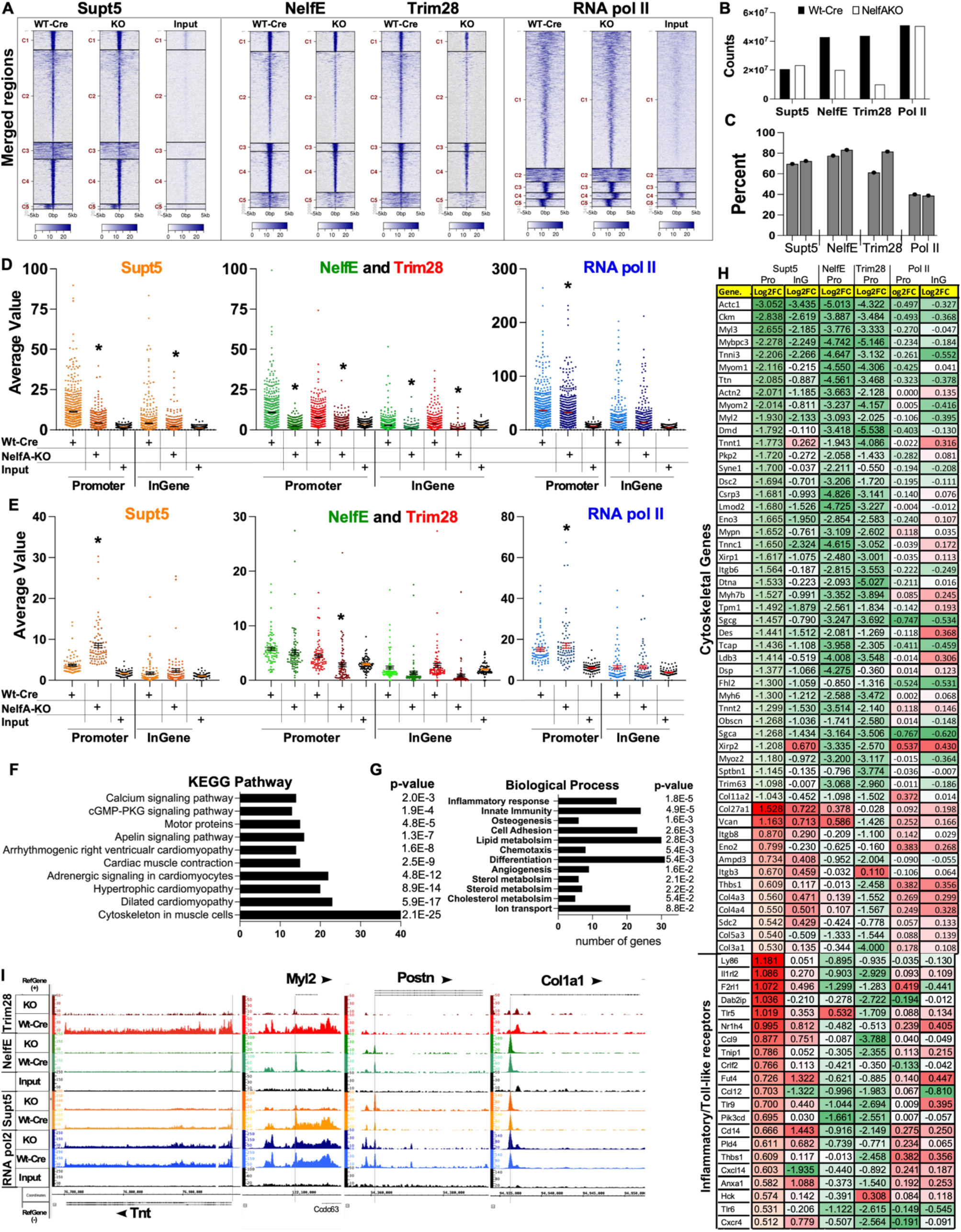
ChIP-seq data from NelfAKO and Wt-Cre mice hearts. **A**. Heatmap shows tag distribution for NelfA complex proteins–Supt5, NelfE, Trim28, and RNA pol II across merged regions (values in z-axis/color, active regions in y-axis) in Wt-Cre and NelfAKO hearts. The data is presented in five clusters (default), labeled C1 to C5, and sorted. **B**. The graph shows the sequencing tag counts for the NelfA complex proteins as indicated. **C**. The graph illustrates the promoter occupancy status of the NelfA complex proteins, as indicated. **D**. Violin plots show the genomic binding status at the promoter and in-gene regions for NelfA complex proteins as indicated, for genes that showed a decrease in Supt5 promoter occupancy. **E**. Violin plots show the genomic binding status at the promoter and in-gene regions of NelfA complex proteins, as indicated, for genes that showed an increase in Supt5 promoter occupancy. Error bars represent SEM. * indicates p < 0.05 compared to the respective Wt-Cre group. **F**. Graphs show the functional annotation of the genes that show a decrease in promoter Supt5, as categorized by KEGG Pathway using DAVID. **G**. Graphs show the functional annotation of the genes that show an increase in promoter Supt5, as categorized by Biological Process using DAVID. **H**. Table lists the genes that were categorized as cytoskeletal genes and inflammatory/TLR pathway genes, along with their change in occupancy status (Log2FC) at the promoter and in-gene regions for the NelfA complex proteins (Supt5, NelfE, Trim28, and RNA pol II). **I**. Screenshot of Integrated Genome Browser (IGB) showing the genomic occupancy of NelfA complex proteins for selected genes in Wt-Cre and NelfAKO hearts, as indicated. The X-axis represents chromosomal coordinates, reference sequence, and gene structure. The Y-axis represents the peak value of genomic occupancy in the signal tracks of the samples (Wt-Cre, NelfAKO, and Input).

KEGG pathway analysis of the genes that show a decrease in Supt5 (Log2FC of 0.5) are mostly categorized as Cytoskeletal genes in muscle cells, involved in cardiac contraction, calcium signaling, and cardiomyopathy (**Fig. 3F**). On the other hand, genes that show an increase in Supt5 (Log2FC of 0.5) are those involved in inflammation, innate immunity, and differentiation (**Fig. 3G**), suggesting that these genes are involved in extracellular signaling, leading to or due to infiltration of immune cells and fibroblast activation. Fig. 3H lists the genes that were categorized as cytoskeleton in muscle cells, extracellular signaling, and inflammatory genes (**Fig. 3H**). The ChIP-seq data highlight the significance of NelfA occupancy in gene transcription, showing that the loss of NelfA disrupts promoter complex assembly at mostly cardiac-enriched genes and hampers gene expression in these hearts.

### Disruption of the NelfA-dependent complex results in transcriptional inhibition of cardiac-enriched cytoskeletal and metabolic genes

Furthermore, to investigate the impact of the disrupted NelfA-dependent complex on gene transcription, we performed total RNA sequencing in Wt-Cre and NelfAKO hearts. We observe significant differential regulation in 465 genes with adjusted p-value ≤ 0.05 (blue and red dots in **Fig. 4A**). We further sorted the genes with Log2FC equal to or greater than 1 (2-fold change, red dots in Fig 4A), which included 123 genes showing downregulation, while 137 genes are upregulated in NelfKO hearts compared to Wt-Cre (**Fig 4A**). KEGG pathway analysis categorized most of the differentially expressed genes (p≤0.05) into those involved in cytoskeletal and metabolic pathways (**Fig. 4B** and **4C**). Interestingly, most of the downregulated genes were identified as those involved in muscle contraction, sarcomere organization, fatty acid metabolic process, transmembrane transport, and negative regulation of apoptosis (**Fig. 4D** and **4G**). Further categorization into biological processes listed transport genes at the top, with 9 downregulated genes involved in potassium transport (**Fig. 4E** and **4F**). On the other hand, genes that show an increase in expression are mostly involved in cellular response to external stimuli and stress, apoptosis and programmed cell death, and senescence (**Fig. 4H**). We predict that the increase in these genes is secondary to the ensuing pathogenesis, rather than directly resulting from the loss of cardiomyocyte NelfA, and may be a contribution from infiltrating immune cells and fibroblasts. Next, we examine the expression of selected genes in NelfAKO and Wt-Cre hearts at 21-25days (no dysfunction on echocardiography) and 55-60days (onset of failure) after birth, which shows that downregulation of contractile genes precedes the cardiac dysfunction and ventricular dilatation (**Fig. 4I** and **4J**).

**Fig. 4.**
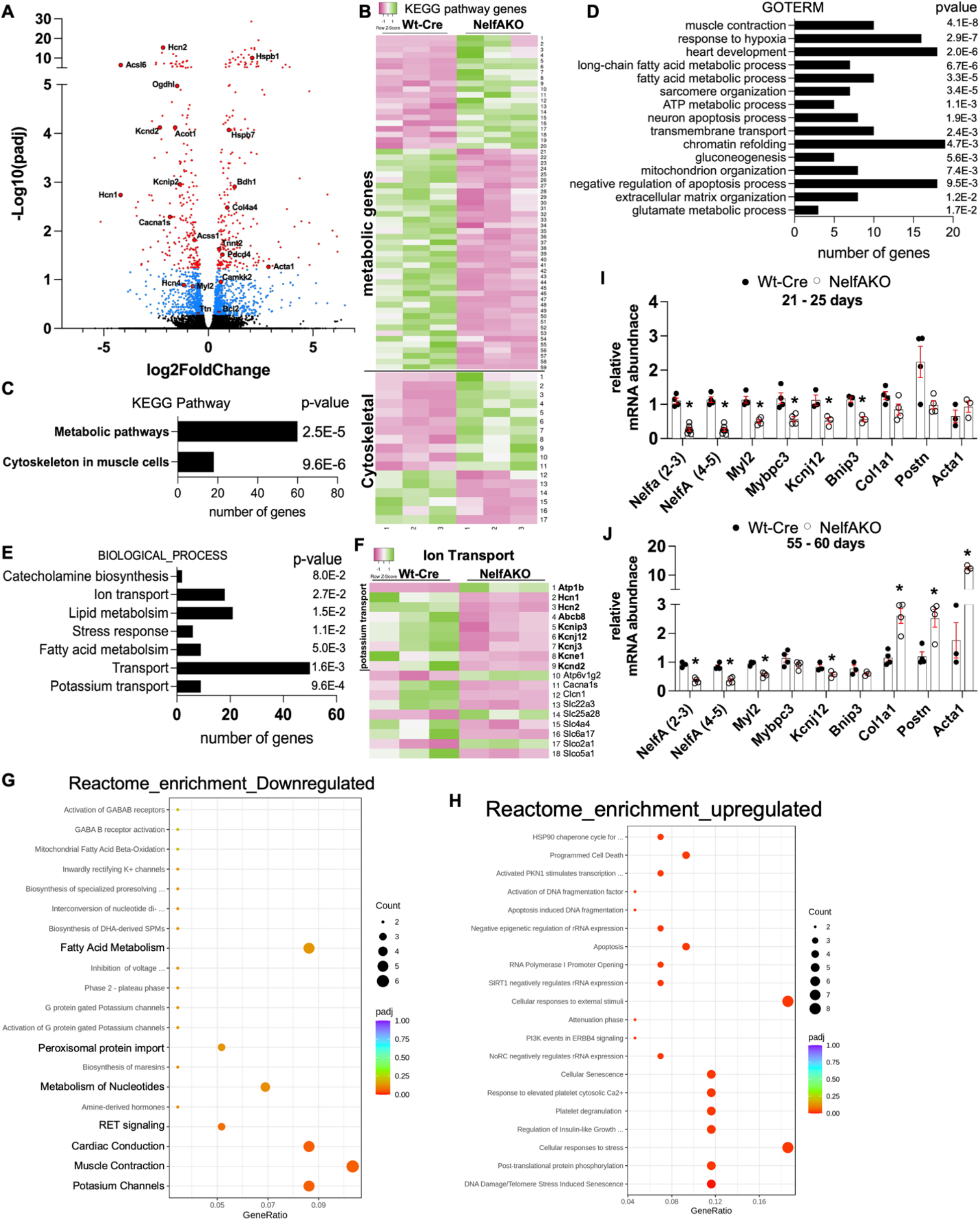
RNAseq data from NelfAKO and Wt-Cre mice hearts. **A**. Volcano plot shows the change in expression of genes between Wt-Cre and NelfAKO hearts as determined by RNAseq analysis. Red dots show genes with Log2FC of 1 and adjusted p<0.05, and blue dots are genes with adjusted p<0.05. **B**. Heatmap shows genes that are differentially expressed between Wt-Cre and NelfAKO hearts (adjusted p<0.05). **C**. The graph shows the number of differentially expressed genes (DEGs) in the indicated functional groups as categorized by KEGG Pathway using DAVID. **D** and **E**. The graph shows the number of downregulated DEGs in the indicated functional groups as categorized by Gene Ontology (GO) TERM and Biological Process, respectively, using DAVID. **F**. Heatmap shows the downregulated DEGs from (E), involved in ion transport. **G** and **H**. Dot plots show the top 20 enriched pathways from Reactome pathway enrichment analysis of DEGs that are decreased or increased, respectively, as indicated. **I** and **J**. The graphs represent mRNA abundance of selected genes, as measured by qPCR, normalized to 18S, in NelfAKO and Wt-Cre hearts from two age groups, 21-25days postnatal and 55-60days postnatal, respectively. For all graphs, error bars represent SEM, * is p < 0.05 compared to respective Wt-Cre, n=3-4 for each group.

### Exogenous NelfA rescues the NelfA-KO phenotype

To determine whether the phenotype observed in the NelfA-KO hearts is a direct effect due to loss of NelfA function, we delivered AAV9-mediated exogenous NelfA in 30-day-old mice (before the onset of cardiac dysfunction) and performed functional and molecular analysis when these mice were 65 days old (**Fig. 5A**). Interestingly, exogenous NelfA restored the nuclear expression levels and the loss of chromatin-bound NelfA in KO hearts (**Fig. 5B** and **5G**). The decrease in cardiac-enriched Myl2 expression was also normalized to control levels in these hearts compared to those in KO hearts (**Fig. 5B**). Further, we observe rescue in heart weight (**Fig. 5C**), ejection fraction (**Fig. 5D**), left ventricular dimensions (**Fig. 5E**), and residual ventricular volume (**Fig. 5F**) in the AAV9-NelfA-injected hearts compared to NelfA-KO hearts. These results validate the cardiomyopathy phenotype developed in mice with the loss of NelfA function in cardiomyocytes.

**Fig. 5.**
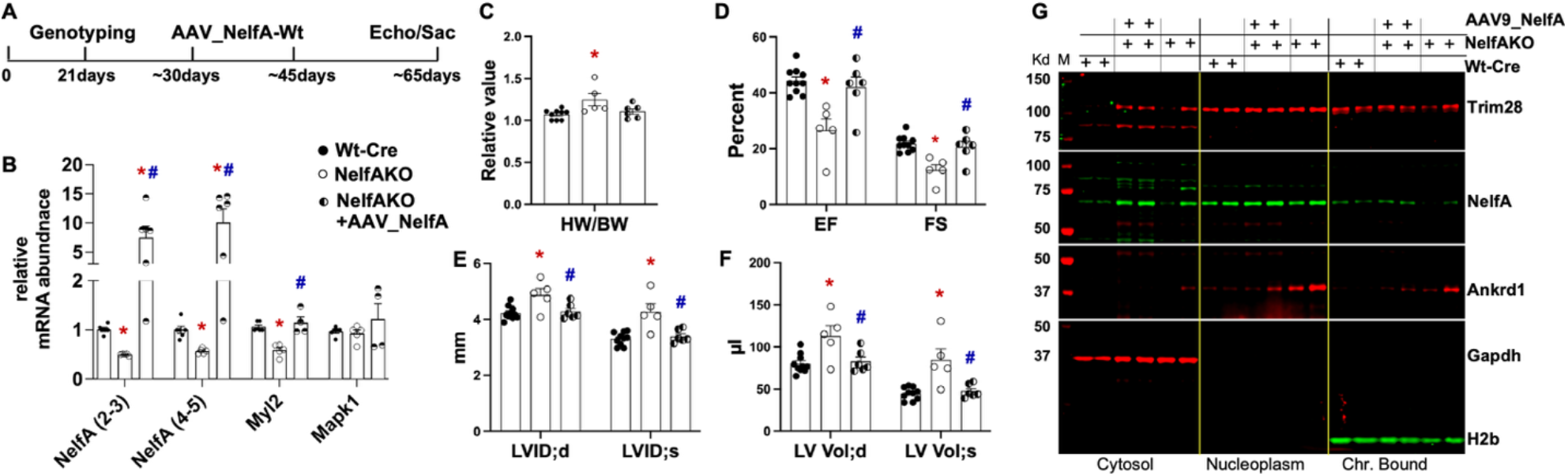
Exogenous NelfA rescues NelfAKO cardiac phenotype. **A**. Schematic shows the AAV9-NelfA-mediated rescue strategy in NelfAKO and Wt-Cre mice. **B**. The graph shows the relative mRNA abundance of selected genes in Wt-Cre, NelfAKO, and NelfAKO + AAV-NelfA hearts. **C–E**. The graphs represent heart weight per body weight (HW/BW), percent ejection fraction (%EF), and percent fractional shortening (%FS), and left ventricular internal dimensions (LVID;s, LVID;d) and volume (LV Vol;s and LV Vol;d) during systole and diastole, respectively, in Wt-Cre, NelfAKO, and NelfAKO + AAV-NelfA mice hearts, as indicated and measured by 2D echocardiography. F. Western blot shows expression levels for indicated NelfA complex proteins, along with their subcellular localization in Wt-Cre, NelfAKO, and NelfAKO + AAV-NelfA mice hearts. Ankrd1 (Carp) is shown as a marker for cardiac hypertrophy. Gapdh and H2B are shown for compartment specificity and as loading controls. For all graphs, error bars represent SEM, * is p < 0.05 compared to respective Wt-Cre, and # is p<0.05 compared to NelfAKO, n=3-9 for each group.

## Discussion

Wolf-Hirschhorn Syndrome is a contiguous gene syndrome, first described in the 1960s, with clinical manifestations ranging from mild to severe outcomes, depending on the extent of deletion in the short arm of chromosome 4 [19]. Characterization of several patients with varying degrees of clinical severity, which includes intellectual and developmental defects, ranging from severe growth retardation, seizures, and classical ‘Greek helmet appearance’, along with midline closure defects, skeletal, and heart malformations, to mild phenotypes that remain underdiagnosed, have led to the identification of two critical regions WHSCR1 and WHSCR2 encompassing several genes [19] [20-22]. WHSC1, a global histone methyltransferase with specificity for Histone 3 at lysine 36 (H3K36me^2^), and WHSC2, *also known as* NelfA, were identified as core genes of the WHSCR, with haploinsufficiency in WHSC1 observed in almost all patients. Furthermore, additional genes, such as LETM1, FGFRL1, and SLBP, were identified and included as part of the critical regions. Interestingly, individual characterization of these genes confirmed the true contiguous and complex nature of WHS and clarified the diverse range of clinical manifestations seen in patients with large deletions versus microdeletions or translocations in these regions [23]. Molecular examination of a panel of cell lines derived from WHS patients shows that a decrease in NELFA and Stem-loop binding protein (SLBP) expression is associated with altered chromatin remodeling, defects in histone deposition, high Histone 3 levels, and an increase in heterochromatin marker, H3K9me3 [24].

Chromatin accessibility and promoter activity regulate gene transcription during development and disease. Adaptation of gene expression is the most fundamental response of the heart to hypertrophic stress [25], with peak increase in RNA pol II activity and total RNA content within 24-48 hours in rats subjected to pressure overload-induced cardiac hypertrophy [26-28]. While examining active gene transcription in hearts, we measured RNA pol II genome occupancy and reported modes of gene transcription in hearts undergoing postnatal growth and pressure overload-induced cardiac hypertrophy. Our data showed that the majority of genes, primarily essential genes, were regulated by the promoter clearance of paused RNA polymerase II, exhibiting incremental changes in expression with cardiac growth [11]. RNA pol II pausing as a mode of eukaryotic transcription regulation was first described in Drosophila, where it was shown to regulate the expression of heat shock genes. Subsequent studies reported widespread RNA polymerase II (RNA pol II) pausing at gene promoters, which plays a central role in gene regulation during development and inflammatory responses. Transcriptional studies have implicated two complexes in mediating pol II pausing: the Negative Elongation Factor (NELF) and the *Drb* sensitivity-inducing factor (DSIF) complex, which is released by P-TEFb (Cyclin T/Cdk9)-dependent phosphorylation [3, 4, 29]. Nelf complex consists of four of the five Nelf proteins, including NelfA (Whsc2), NelfB (Cobra), NelfC or D (TH1L), and NelfE (RDBP). Structural examination of the paused complex establishes the orientation of Nelf and Supt5 associations with pol II, and identifies novel aspects of the complex like, 1) positive amino acid residues patches on NelfAC subcomplex, 2) Four such positive patches exist in NelfA-NelfC subcomplex, facing opposite to its association with NelfB, and can bind to single-stranded nucleic acids, 3) NelfA is indispensable for the formation of the paused complex, and 4) two tentacle-like extensions from NelfA and NelfE that bind to RNA pol II and nascent transcript, respectively [9, 10].

In this study, we characterize the NelfA-dependent complex in mice with postnatal loss of NelfA function, focusing on the assembly of the complex, gene expression, and its functional outcomes. Previous studies have proposed Nelf-mediated RNA Pol II pausing as a mechanism to maintain a nucleosome-free promoter for continuous gene transcription [30], which agrees with our results showing the highest NelfA peaks in constitutively expressed essential genes in quiescent and hypertrophied hearts [13]. Nelf-dependent Pol II pausing has been shown to be a prerequisite for transcript processing, such as capping and splicing [31]. Our interactome data from sham and TAC-operated hearts reveal interactions of chromatin-bound NelfA with Trim28 (<u>tri</u>partite <u>m</u>otif-containing protein <u>28</u>) and Adprh1l in the heart, along with other Nelf proteins, RNA polymerase II subunits, and Supt5 (DSIF subunit). Trim28, *also known as* KAP1 (<u>K</u>RAB-<u>a</u>ssociated <u>p</u>rotein <u>1</u>) or TIF1β (transcription intermediary factor 1 beta), has been shown to regulate the pausing and release of RNA polymerase II (pol II), which is dependent on the Trim28 phosphorylation status [32]. Trim28, while dubbed as a transcriptional repressor, based on domain interaction and in vitro studies, has been associated with epigenetic modifications and structural chromatin changes. Recently, Trim28 has been shown to control the entry and Cdk9-dependent exit of RNA polymerase II (RNA pol II) from the paused state in cancer cells, thereby sustaining the expression of specific genes [33]. Our RIME analysis reveals that Trim28 and Supt5, which have been shown to associate with the non-template strand [32] [34], are integral components of the NelfA-dependent complex in heart tissue, under both quiescent and hypertrophic conditions, and may be involved in regulating active gene transcription. Supt5 (suppressor of Ty5, homolog), a component of DSIF, associates with pol II, and is a prerequisite for Nelf binding to pol II-Supt5 subcomplex [3, 35]. P-TEFb-dependent phosphorylation of DSIF can switch its function from a transcription repressor to an activator of productive elongation [36]. KOW4-5 regions of Spt5 have been shown to mediate pol II pausing [37]. In addition to RNA pol II and other Nelf proteins, Adprhl1 also precipitated as part of the chromatin-bound NelfA in our RIME data. Few studies have characterized this protein, reporting it as enriched in the heart and required for cardiac development and chamber specification [38, 39]. Loss of Adprhl1-ROCK signaling in human embryonic stem cell line H9 resulted in defects in adhesion and disrupted electrical conduction and calcium transients [40]. Our occupancy data with selected associating proteins in KO hearts show a significant decrease in genome occupancy of these proteins, suggesting hampered assembly of the complex at cardiac-enriched gene promoters. The transcriptome data in these hearts correlate with the ChIP-seq data, which show a significant decrease in the expression of cytoskeletal and metabolic genes. Interestingly, we observe a decrease in potassium channel genes, which could have a detrimental effect on cardiomyocyte electrophysiological conduction and contraction. We believe that the contractile dysfunction leads to cardiomyopathy, and the differential regulation of the metabolic genes could be secondary and further contribute to the development of heart failure and death.

In conclusion, the NelfA-dependent transcriptional complex, comprising components involved in active transcription, chromatin remodeling, and pre-mRNA processing, plays a crucial role in the expression of cardiac-enriched genes and is essential for maintaining postnatal cardiomyocyte contractile and metabolic homeostasis, as well as for cardiac development and function.

## Declaration of interest

The authors have no conflicts of interest with the contents of this article.

## Funding

This work was supported by National Heart, Lung and Blood Institute (NHLBI) of National Institute of Health (NIH) funding to the corresponding author (R01HL150059).

## Author contributions

SA performed the experiments; CT and AV assisted with the genotyping, experimentation, microscopy, and histology of the sections; ZY performed the sham/TAC surgeries, as well as the echocardiography on the mice; AI performed the histology and staining; DS designed experiments, performed data analysis with figures, and wrote the manuscript.

## Acknowledgements

We thank Dr. Sadoshima’s lab for providing us with the αMHC-Cre mice; permission was also obtained from Dr. Molkentin, Director of the Division of Molecular Cardiovascular Biology at Cincinnati Children’s, via email. We also thank Dr. Sadoshima, Chair of the Department of Cell Biology and Molecular Medicine, Rutgers New Jersey Medical School, for his support.

## Notes

### Competing Interest Statement

The authors have declared no competing interest.

